# Propofol Mediated Unconsciousness Disrupts Progression of Sensory Signals through the Cortical Hierarchy

**DOI:** 10.1101/2023.06.25.546463

**Authors:** John M. Tauber, Scott L. Brincat, Emily P. Stephen, Jacob A. Donaghue, Leo Kozachkov, Emery N. Brown, Earl K. Miller

## Abstract

A critical component of anesthesia is the loss sensory perception. Propofol is the most widely used drug for general anesthesia, but the neural mechanisms of how and when it disrupts sensory processing are not fully understood. We analyzed local field potential (LFP) and spiking recorded from Utah arrays in auditory cortex, associative cortex, and cognitive cortex of non-human primates before and during propofol mediated unconsciousness. Sensory stimuli elicited robust and decodable stimulus responses and triggered periods of stimulus-induced coherence between brain areas in the LFP of awake animals. By contrast, propofol mediated unconsciousness eliminated stimulus-induced coherence and drastically weakened stimulus responses and information in all brain areas except for auditory cortex, where responses and information persisted. However, we found stimuli occurring during spiking Up states triggered weaker spiking responses than in awake animals in auditory cortex, and little or no spiking responses in higher order areas. These results suggest that propofol’s effect on sensory processing is not just due to asynchronous down states. Rather, both Down states and Up states reflect disrupted dynamics.

## Introduction

General anesthesia is used in nearly 60,000 procedures every day in the United States (Brown et al., 2010). A critical component of general anesthesia is unconsciousness, during which a patient is unaware of their environment (Brown et al., 2010). Intraoperative awareness occurs when this goal is not achieved (Ghoneim, 2000). Although the phenomenon is rare (Sebel et al., 2004), patients that experience it report severe trauma (Kotsovolis & Komninos, 2009). Most studies of anesthetic effects on the brain have focused on physiological state change. But if we are to understand how anesthesia renders unconsciousness and how this fails in intraoperative awareness, we need to understand its effects on processing of sensory inputs. We aimed to do so using propofol, one of the most commonly used anesthetics.

Propofol is a GABA-agonist (Bai et al., 1999; Hemmings et al., 2005, 2019). Although propofol’s molecular mechanism of action is well understood (Sahinovic et al., 2018), we have less understanding of how it works at the level of functioning networks (Purdon et al., 2013; Brown et al., 2011; Lewis et al., 2012, 2013). Propofol induces overall increases in slow oscillations (0.1-4Hz) in electroencephalogram (EEG) and local field potential (LFP) recordings, and a broad reduction in spiking activity (Bastos et al., 2021; Redinbaugh et al., 2020; Purdon et al., 2015). Spiking becomes strongly coupled to the phase of the slow oscillations, creating alternating irregular “Up” and “Down” states of high and low activity, respectively (Lewis et al., 2012; Bastos et al., 2021). This may disrupt long-range synchronization between cortical regions, a putative mechanism for cortical communication (Crowe et al, 2013; Lewis et al., 2012; Bastos et al., 2021; Redinbaugh et al., 2020; Fries, 2015; Pesaran et al., 2018). Support for this comes from observations that sensory responses are weakened in higher cortex but can be preserved in lower (sensory) cortex (Krom et al., 2020; Ishizawa et al., 2016). However, few studies, especially in animals, have followed the chain of sensory processing from lower to higher cortex.

We did so using data collected in two non-human primates from our previous study on propofol anesthesia (Bastos et al., 2021). We compared and contrasted cortical responses to auditory and tactile stimulation before and after loss of consciousness. This analysis includes simultaneous recordings of LFP and spiking activity from multiple cortical levels: sensory (temporal) cortex, associative (parietal) cortex, and higher (frontal) cortex. Our results suggest propofol anesthesia leaves intact sensory processing in the sensory cortex but these signals fail to be transmitted to higher-level cortical areas.

## Materials and Methods

### Experimental

Two non-human primates (rhesus macaques - Macaca mulatta, abbreviated NHP) participated in the study (Bastos et al., 2021). NHP 1 was male, aged 14 years and weighed 13.0kg. NHP 2 was female, aged 7 years and weighed 5.0kg. A total of 21 sessions (11 from NHP 1, 10 from NHP 2) were used. Each session consisted of 15-90 min of awake baseline electrophysiological recordings. Then propofol was intravenously infused via a computer-controlled syringe pump. To induce unconsciousness a high-rate infusion was given for 30 minutes (285 *µ*g/kg/min for NHP 1; 580 *µ*g/kg/min for NHP 2). Immediately following the high-rate infusion was 30 minutes of a maintenance dose (142.5 *µ*g/kg/min for NHP 1; 320 *µ*g/kg/min for NHP 2). The infusion rates were determined based on the animals’ age and weight. Infusion was performed via a subcutaneous vascular access port at the cervicothoracic junction of the neck with the catheter tip reaching the termination of the superior vena cava via the external jugular vein.

Both NHPs were chronically implanted with four 8 x 8 iridium-oxide contact microelectrode arrays (‘Utah arrays’, MultiPort: 1.0 mm shank length, 400 *µ*m spacing, Blackrock Microsystems, Salt Lake City, UT), for a total of 256 electrodes. Arrays were implanted in the prefrontal (area 46 ventral and 8A), posterior parietal (area 7A/7B), and temporal-auditory (caudal parabelt area STG [superior temporal gyrus]) cortices. We refer to the recorded brain areas in the text as Superior Temporal Gyrus (STG), Posterior Parietal Cortex (PPC), Region 8A (8A), and Prefrontal Cortex (PFC) (Figure 1A).

**Figure 1:**
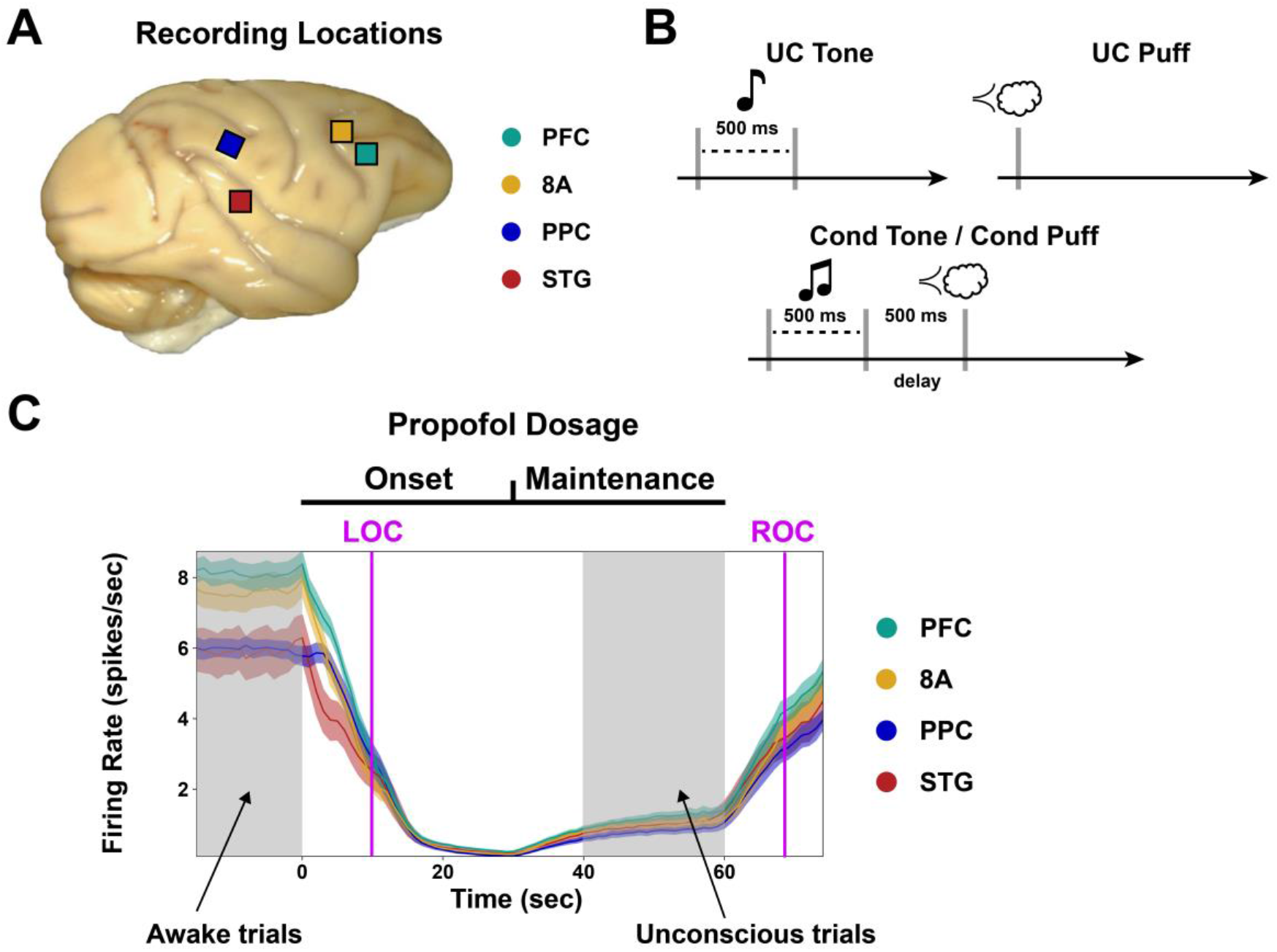
LFP and spiking data were recorded from four cortical areas while stimuli were delivered in Awake and Unconscious animals. (A) Implanted locations of Utah arrays. STG: Superior Temporal Gyrus (Auditory); PPC: Posterior Parietal Cortex (Associative); 8A: Region 8A (Cognitive); PFC: Prefrontal Cortex (Cognitive). (B) Stimuli delivered during Awake and Unconscious states. UC (unconditioned) Tone and Cond. Tone are two distinct sounds, both lasting half a second. Cond. Tone is followed by an air puff delivered to the animals’ face following a half-second delay. UC Puff is an identical air puff without any preceding tone. (C) Spike rates are stable during Maintenance dose. Average firing rates (1-minute bins) displayed for each brain area (mean +/SEM). Horizontal black bars (top) indicate time course of propofol infusion dosage. A high dose was given for 30 minutes (Onset) followed by 30 minutes of a lower dose (Maintenance). Purple bars mark average time for Loss of Consciousness (LOC) and Return of Consciousness (ROC). Time ranges used for Awake and Unconscious states indicated by gray boxes.

LFPs were recorded at 30 kHz and filtered online via a lowpass 250 Hz software filter and downsampled to 1 kHz. Spiking activity was recorded by sampling the raw analog signal at 30 kHz, bandpass filtered from 250 Hz to 5 kHz, and manually thresholding. Blackrock Cereplex E headstages were utilized for digital recording via 2-3 synchronized Blackrock Cerebus Digital Acquisition systems. Single units were sorted manually offline using principal component analysis with commercially available software (Offline Sorter v4, Plexon Inc, Dallas, TX). All other pre-processing and analyses were performed with Matlab (The Mathworks, Inc, Natick, MA).

Three different stimuli were presented to the NHPs throughout each session to characterize sensory processing under anesthesia (Figure 1B). Sensory stimuli were delivered continuously throughout the experimental sessions. They were spaced approximately five seconds apart and randomly interleaved. Each stimulus was presented roughly 60-80 times in both Awake and Unconscious states for each session, providing samples sizes comparable to previous studies (Purdon et al., 2013; Ishizawa et al., 2016).

The number of sorted units per brain area differed between NHP 1 and NHP 2, and also differed slightly across sessions. For NHP 1: ∼50 units were recorded in STG, ∼80 in PPC, ∼40 in 8A, and ∼40 in PFC. For NHP 2: 1-5 units were recorded in STG, ∼20 in PPC, ∼20 in 8A, and ∼80 in PFC. We excluded areas STG, PPC, and 8A in NHP 2 from the analysis of spiking responses. In the STG of NHP 2, we were unable to resolve Up and Down states due to the low number of neurons. Up states were observed in the PPC and 8A of NHP 2, but had very short duration, and hence too few stimuli occurred within them to estimate a response with high confidence. For further information on neural recording and propofol delivery methods see (Bastos et al., 2021). All procedures followed the guidelines of the MIT Animal Care and Use Committee (protocol number 0619-035-22) and the US National Institutes of Health.

We analyzed the cortical responses to sensory stimuli under two different conditions: Awake (before administration of propofol) vs Unconscious (after loss of consciousness, during the maintenance dose). Data from the maintenance dose was used for two reasons. First, the brain state was unchanging during the maintenance dose (Figure 1C). This consistency allowed for more reliable data pooling across stimulus presentations and comparison with the pre-anesthesia awake state, which was also consistent over time. Second, the maintenance dose was designed to mimic the portion of human clinical procedures when there is also steady-state maintenance. Thus, understanding how propofol alters cortical sensory processing has the most clinical relevance during this interval. We used data from both NHPs in our analyses. All results were consistent across NHPs unless noted otherwise.

### Data Analysis

#### Spectral Analysis

All spectrograms and coherograms were computed using 400 ms length windows with 50 ms step sizes. We used multitaper spectral estimation to improve power spectral density estimation (Babadi & Brown, 2014). The number of tapers were selected using floor(2TW-1) tapers, where T is the length of the window in seconds and W is the half-bandwidth of our desired spectral resolution. We used W=4, which resulted in two tapers.

Average spectrograms for each brain area were computed across electrodes and trials for a given session. Reported averages were then calculated across sessions. To display the change in oscillatory activity related to stimulus presentation, baseline power was calculated using 500-200 ms before the stimulus was given. The change from baseline was then calculated as:

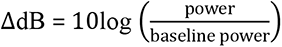

Coherence was calculated as described in (Kramer & Eden, 2016). The average coherence reported for each brain area pair was found by taking the average coherence across all pairs of electrodes from the two areas. Baseline normalization was performed by subtracting the average baseline coherence value (500- 200 ms before stimulus). Non-phase locked power and coherence were calculated by subtracting the trial-averaged time domain response from each trial prior to computing the frequency response.

#### Stimulus Decoding

Stimulus trial data was first binned in 20 ms intervals. For each 20 ms bin, the population data was ‘flattened’ by concatenating timepoints. That is, a matrix of size (electrodes, time) would be transformed to an array of length (electrodes x time). This method was chosen to retain all available information. For each bin of data, we used a linear support vector classifier (SVC in Python’s sklearn package (Pedregosa et al., 2011), with default regularization parameter C=1) for decoding analysis (Hastie et al., 2009). Decoding analysis was performed separately on each experimental session. For a given session, a decoder was trained on labeled data (UC Tone, UC Puff, or Cond Tone) from a given time bin. Accuracy was then assessed on held out test data. Average test accuracy from a 10-fold cross-validation was calculated for each session. (That is, we held out a test set containing 10 percent of trials and trained on the remaining 90 percent of the data. The session test accuracy was computed by repeating this process 10 non-overlapping test sets that partitioned the data and taking the average test accuracy. The final means and standard errors were then calculated as the average test accuracy across sessions.

#### Statistical Analysis

Statistical tests for changes in coherence were performed by first transforming the values to be normally distributed and then using standard t-tests (Jarvis and Mitra, 2001). Non-parametric tests (Wilcoxon) were used for all other statistical comparisons (one-sample for stimulus vs baseline, two-sample for Awake vs. Unconscious). Multiple comparison corrections were performed using False-Discovery Rate (FDR) correction with a family-wide error rate of 0.01 (Seabold and Perktold, 2010).

### Hidden Markov Model for Estimating Spiking Responses

#### Model Specification

To account for the presence of Up and Down states in the Unconscious state, we designed an HMM to estimate spiking stimulus responses (summarized in Figure 2). Before introducing the formal structure of the model, we provide a brief conceptual description. Each neuron has its own mean firing rate parameters, one for Up states and one for Down states (Figure 2A-D). In the absence of a stimulus, each neuron’s firing rate is a function of the mean rate parameters, the hidden state, and the neuron’s spiking history (Figure 2C). Finally, the stimulus response is modeled using a spline basis and separate sets of coefficients depending on which state *stimulus onset* occurs in (Figure 2C,D). The parameters governing the stimulus response are shared across neurons such that we estimate an average population response for each area.

**Figure 2:**
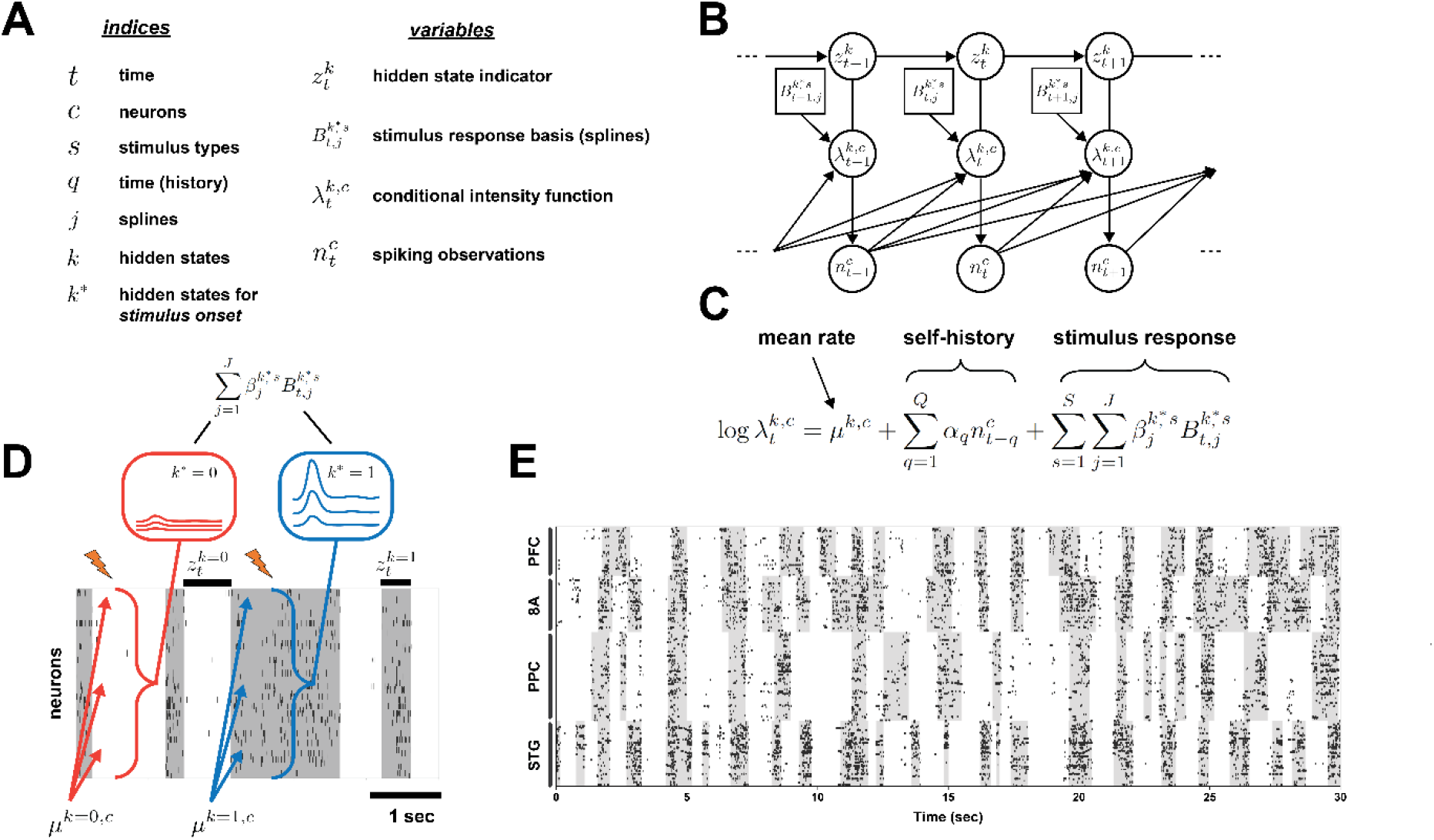
Hidden Markov Model used for estimating spiking stimulus responses in the Unconscious state. (A) Model notation. (B) Graphical model structure of the HMM. (C) The Conditional Intensity Function (CIF) for a neuron in the model. Each neuron has a distinct mean rate parameter that differs between Up and Down states. Self-history and stimulus response parameters are shared across neurons. (D) Stimulus response schematic. In the Down state, neurons hypothetically will have a low firing rate (red arrows) and minimally respond to the stimulus (red box). In Up states, baseline firing rates increase (blue arrows) and stimulus response is heightened (blue box). (E) Raster plot showing 30 seconds of spiking data from NHP 1 during the Unconscious state. Up and Down spiking states are visible in each brain area. Gray boxes indicate Up state labels produced by HMM algorithm.

Let *k* ∈ {0, …, *K*} be the discrete states in the system. In the analysis of spiking responses in the Unconscious state, we let *K* = 2 with *k* = 1 and *k* = 0 representing the Up and Down states, respectively. Next, we define an indicator function *z_t_*^*k*^ which equals 1 if the system is in state *k* and 0 otherwise. Let ***z***_***t***_ = (*z_t_*^1^, …, *z_t_*^*K*^) refer to the vector of indicator functions for all hidden states. The state transition probabilities are represented in a state transition matrix:

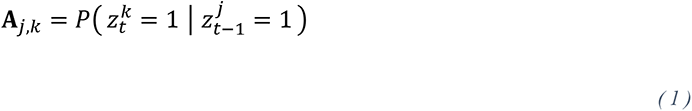

which is treated as an unknown parameter to be estimated. We use ***π*** to denote the initial state probabilities. The spiking observations for neuron *c* ∈ {1, …, *C* } are modeled as a point process (Daley et al., 2003). Letting *N*_(*a*,*b*]_^*c*^ denote the number of spikes occurring in the time range (*a*, *b*] for *a* < *b* and *H_t_*^*c*^ denote the history of the process up to time *t*, the Conditional Intensity Function (CIF) for neuron *c* is defined as:

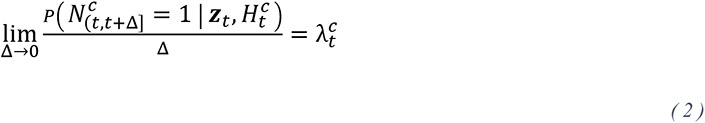

where Δ is the time-bin width and 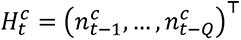 is the neurons self-history for time lag *q* ∈ {1, …, *Q*}. We assume that for a small Δ (typically 1 millisecond as in our study) we can make an accurate discrete-time approximation to the continuous-time CIF at time *t* as

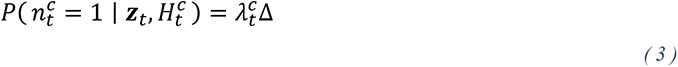

where λ*_t_*^*c*^Δ is linked to the hidden states by:

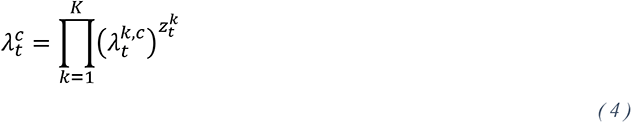

Next, let *s* ∈ {1, …, *S*} represent the different stimulus types (e.g. UC Tone). We define the CIF for each state *k* to be a function of a mean rate, spiking history, and the sensory stimulus:

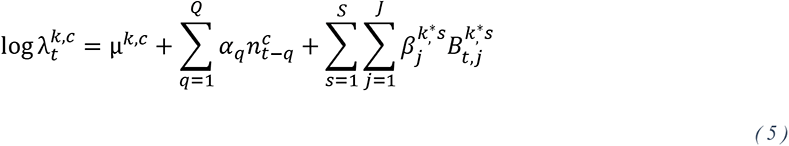

where we have history coefficients (*α*_1_, …, *α*_Q_)^⊤^, *s* ∈ {1, …, *S*} the stimulus type, *k*^∗^ is the hidden state in which *stimulus onset* occurred, and the 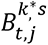 serves as a basis for modeling the stimulus response. We note that the history terms create conditional dependencies across observations over time, which is not standard in HMMs. However, because these dependencies occur in the observation equation, no major changes to the estimation algorithm are required (this structure is commonly seen in ‘auto-regressive HMMs’) (Hamilton, 1990; Murphy, 2023).

We assume a common response across neurons for several reasons: first, as we cannot ensure identical neuron populations are recorded in each session, we target a mean population response that can be easily combined and compared across sessions. Second, limited stimulus response data per session might preclude reliable individual parameter estimates for each neuron. Finally, the computational burden of fitting a model with distinct stimulus responses parameters for each neuron would be substantial. While acknowledging the oversimplification of this biophysical assumption, we argue that a model estimating a mean population response in each area, alongside individual mean firing rates, strikes a balanced compromise between accuracy and practicality, adequately addressing our study’s research questions.

#### EM Algorithm

Given the observed spiking data our objective is to estimate the parameters:

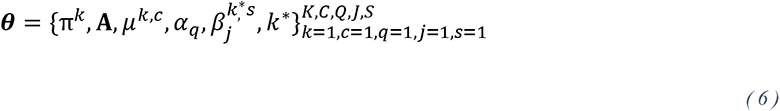

We estimate the parameters and hidden states using the Expectation Maximization (EM) algorithm (Dempster et al., 1977), a standard approach for HMMs and latent variable models more generally (Bishop, 2006; Murphy, 2023). The EM algorithm works by iterating between 1) taking the expected value of the complete-data log likelihood with respect the conditional distribution of the hidden states given the data and 2) maximizing the expected complete-data log likelihood over ***θ*** (Dempster et al., 1977). We define the *Q*- function *Q*(***θ*** ∣ ***θ***^(*r*)^) as the expectation of the complete-data log likelihood, on iteration *r* + 1 of the EM algorithm, using the parameters ***θ***^(*r*)^from the previous iteration.

##### E-Step

In the E-Step the conditional distribution of the hidden states given the observed data is computed. We use the standard approach of the forward-backward algorithm, and refer the reader to (Rabiner, 1989; Bishop, 2006; Murphy, 2023) for details.

##### M-Step

In the M-Step, we maximize ***θ*** in the *Q-*function. The state parameters ***π*** and **A** have standard closed form solutions. Closed form solutions do not exist for the point process parameters and they must be found numerically. We use the BFGS algorithm (Fletcher, 2013; Virtanen et al., 2020) for this purpose. Finally, after updating the other parameters, we run the Viterbi algorithm to determine the most likely state path, which is used to update the *k*^∗^assignments.

In preliminary work we found that the Up and Down state labels produced by our model switched between states too frequently, likely due to the high time resolution and binary nature of the data. We found putting a Dirichlet prior distribution on each row of the **A** matrix to favor self-transitions (i.e. probabilities along the diagonal) improved the labels (Linderman et al., 2020). Specifically, denoting the *j*^th^ row of the matrix ***A*** as ***A***_*j*,1:*K*_, we put a prior distribution on each row:

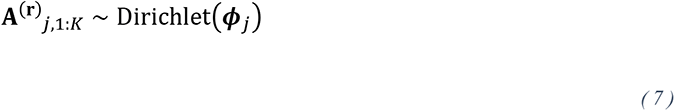

where

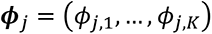

The same parameters values are used for each row, adjusting for the position of the row entry that corresponds to the diagonal element of **A**. Noting that the Dirichlet distribution is the conjugate prior for the Categorical distribution, the closed-form parameter update is (Murphy, 2023):

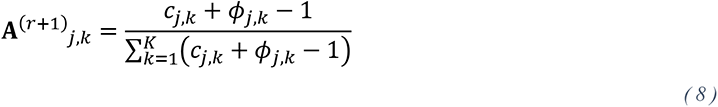

using

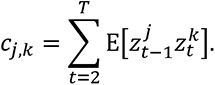

#### Estimated Stimulus Response and Confidence Intervals

For visualization purposes (Figure 6), we defined an estimated stimulus response summary across all neurons for a session:

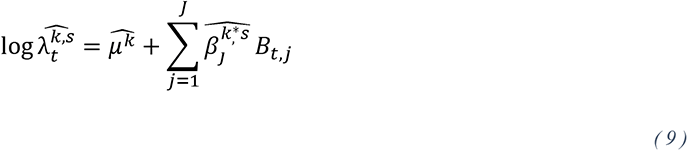

where

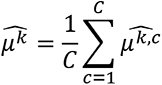

The variance of parameter estimates was calculated using the method of (Louis, 1982), which enables calculating the Observed Fisher Information when using the EM algorithm for producing maximum likelihood estimates (Efron & Hinkley, 1978). On some sessions we found that the variance of small number (fewer than 10 percent of neurons) of the *μ*^*k*,*c*^ parameters were very high. These neurons typically showed very low firing rates and were removed for the calculation of the stimulus response. Confidence intervals were then calculated using the delta method (Cramér, 1999).

While the precise neurons recorded across sessions often vary, we assume the mean firing rate and stimulus response to be relatively consistent and statistically independent across sessions. We therefore combined session-level results and uncertainty by taking an average of the parameter estimates across sessions. Within sessions, the *μ*^*k*,*c*^ were assumed to be independent from each other and from the *β*_*j*_^*k*∗*s*^ parameters. Following the asymptotic normality of maximum likelihood estimators, we assume the parameter estimates from each session are Gaussian distributed and combined estimates across sessions following the properties of summation for Gaussian random variables (Pawitan, 2001).

#### Implementation

We implemented our model as a custom observation class in the SSM package (Linderman et al., 2020). We first applied a Poisson HMM to the data and used the fitted values to initialize ***π***, ***A*** and the *μ*^*k*,*c*^ parameters in our model. The α_q_ and *β*_*j*_^*k*∗*s*^ parameters were initialized at 0. We applied the HMM to spiking data during the Unconscious state, which contained Up and Down states. We used a value of 1 × 10^12^ for the ‘diagonal component’ of ***ϕ***_*j*_ for NHP 1 in all brain areas and 1 × 10^8^ for PFC in NHP 2. The off-diagonal elements of ***ϕ***_*j*_ were set to 1, and the row was then normalized to sum to one (Linderman et al., 2020). A separate model instance was used to estimate the Awake response using *K* = 1 (i.e. constant hidden state). In this case, the EM algorithm is not needed. Rather, the parameter estimation described in the M-Step above is simply performed once.

For self-history we used *Q* = 10 to account for refractory periods. For the stimulus response basis 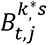 we used natural cubic splines with knots spaced 50 ms apart (Bartels et al., 1995). This basis used *J* = 24 and spanned from 200 ms before stimulus to 1000 ms after stimulus. To ensure that the basis functions were non-overlapping across stimulus types, we started the basis for the Conditioned Puff at stimulus presentation time, resulting in *J* = 20. When plotting the estimated response for the Conditioned Puff, we used the *β*_*j*_^*k*∗*s*^ parameters from the end of the Conditioned Tone estimated response that occupied the 200 ms preceding the Conditioned Puff stimulus.

## Results

We administered propofol to two non-human primates (NHPs). A total of 21 experimental sessions were performed (11 sessions NHP 1, 10 sessions NHP 2). They were first given a high dose of propofol (285- 580 mcg/kg/min) for thirty minutes to induce loss of consciousness. This was followed by thirty minutes of a lower maintenance dose (70-320 mcg/kg/min), similar to that used to maintain anesthesia in human surgeries (see Methods for details about propofol administration). Local field potential (LFP) and spiking were recorded simultaneously from Utah arrays implanted in four cortical areas: Superior Temporal Gyrus (STG), a secondary auditory center; Posterior Parietal Cortex (PPC), an associative region; Region 8A (8A) and Prefrontal Cortex (PFC), cognitive areas (Figure 1A).

Three different stimuli were presented to the NHPs throughout each session to characterize sensory processing under anesthesia. One was a puff of air to the face. As in our previous study, the lack of behavioral response to the airpuff, along with other measures such as eyes opening and closing, was used to determine loss and subsequent recovery of consciousness (see (Bastos et al., 2021)). We also presented two distinct brief (500 ms) auditory tones (Figure 1B). One tone always preceded an airpuff by 500 ms. We call this the Conditioned Tone (Cond Tone). A second tone (Unconditioned Tone, UC Tone) was always presented alone. We also presented airpuffs alone without any preceding tone (Unconditioned Airpuff, UC Puff).

### Anesthesia Alters Cortical Responses to Sensory Stimuli

Cortical responses to stimuli were altered during the Unconscious state relative to the Awake state. We compared LFP spectral power in the Awake vs Unconscious states (Figure 3). During the Awake state, the Cond Tone, Cond Puff, and UC Puff evoked an increase in power (relative to pre-stimulus baseline, see Methods) in the alpha/beta (8-25 Hz) band for all cortical areas (Figure 3B-C, contour lines indicate p < 0.05, Wilcoxon). During the Awake state, the STG also showed a broadband increase in higher-frequency power (< 30 Hz) to all stimuli (Figure 3A-C). The higher frequency response was weaker in the higher cortical areas. The UC Tone elicited weaker responses than the Cond Tone, Cond Puff and the UC Puff (Figure 3A, contour lines indicate p < 0.05, Wilcoxon). During the Unconscious state, the alpha/beta response was decreased in STG and virtually disappeared in the higher cortical areas (Figure 3B-C). In STG (but not other areas), there was an increase in the higher frequency broadband response to the Cond Tone, UC Tone and UC Puff (but not the Cond Puff) relative to the Awake state (Figure 3A-C).

**Figure 3:**
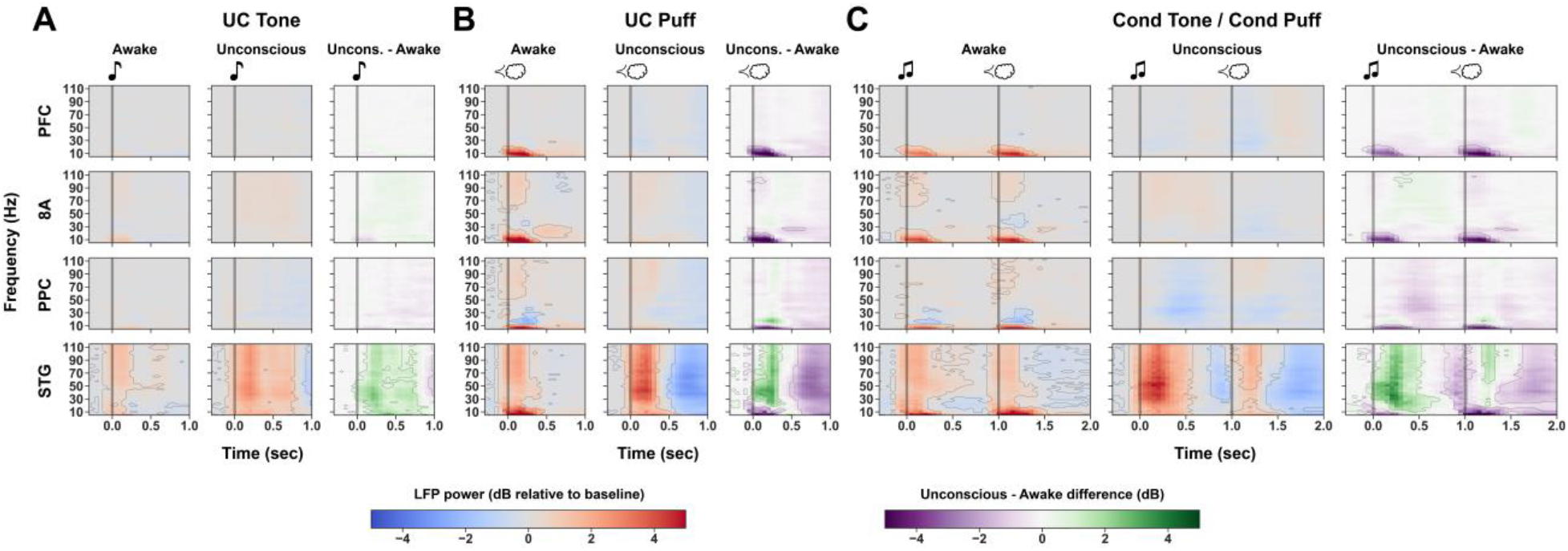
Alpha/beta frequency range responses are lost in higher order cortical areas during anesthesia. (A-C): Change in time- frequency power relative to pre-stimulus baseline in Awake and Unconscious state for each stimulus. Vertical gray bar indicates stimulus onset time. Rightmost panels for each stimulus show the difference in power between Awake and Unconscious responses. Contour lines show regions of statistical significance (Wilcoxon test with FDR correction for multiple comparisons). Stimulus- induced alpha/beta responses disappear during anesthesia, especially in the higher-level cortical areas. Area STG also shows an increase in higher frequency power under propofol.

### Stimulus Information is Progressively Lost through the Cortical Hierarchy During Unconsciousness

Next, we turn to measures of stimulus information. We used a linear classifier to decode stimulus identity using LFP and spiking (see Methods for details). To isolate the activity to individual stimuli, we looked at decoding for stimuli for which there was no immediately preceding stimulus (i.e., UC Tone, Cond Tone, and UC Puff).

During the Awake state, stimulus information was found in all regions of cortex (Figure 4A). During the Unconscious state, stimulus information showed a significant decrease in all areas, as measured by change in the maximum decoding performance achieved during a 50-250 ms post-stimulus time range (Figure 4B; Wilcoxon; *** p < 0.001). In STG, the decrease in stimulus information was relatively modest. Classifier decoding remained well above chance (Figure 4A; Wilcoxon; p < 0.001). This suggests that a significant amount of stimulus information still reaches intermediate-level sensory cortex. By contrast, in the Unconscious state, stimulus information progressively decreased across areas and was near chance levels in PFC, relative to the Awake state.

**Figure 4:**
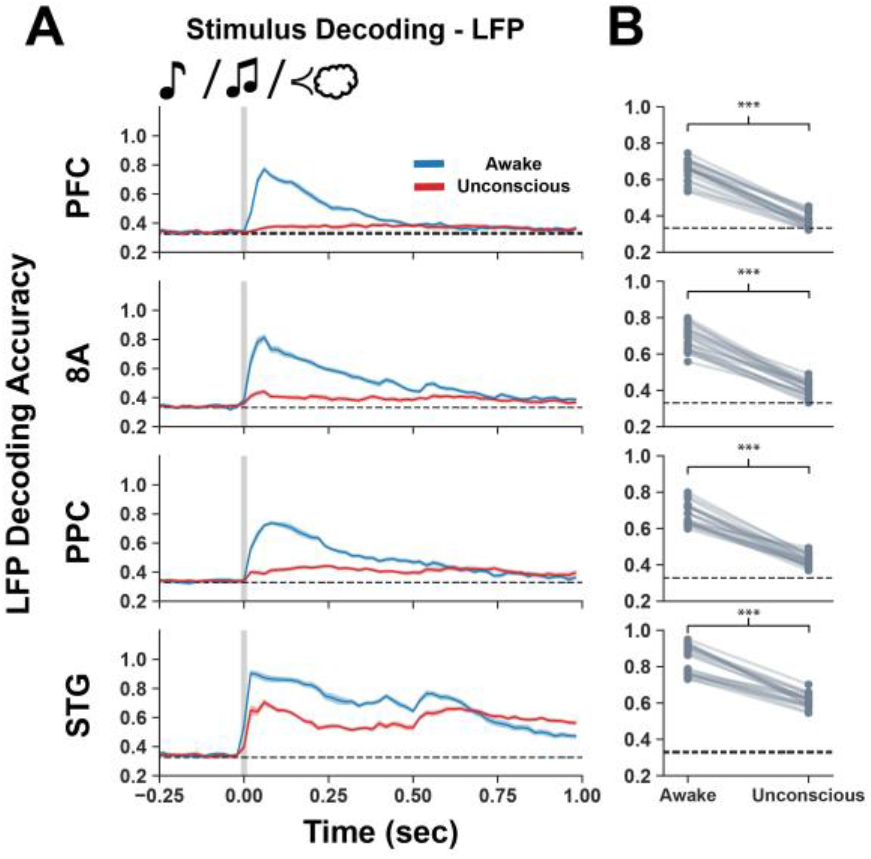
Stimulus information is lost progressively through cortical hierarchy during anesthesia. (A) Decoding accuracy on LFP between the UC Tone, UC Puff, and Cond Tone in each cortical area. Shaded region represents +/- SEM. (B) Change in average decoding accuracy between 50-250 ms post-stimulus for each experimental session. Significance markers indicate difference between Awake and Unconscious (Wilcoxon; p < 0.001).

### Intracortical Coherence Is Disrupted During Anesthesia

Our results suggest that propofol anesthesia disrupts feedforward transmission of sensory information from sensory (e.g., STG) to higher cortical areas (e.g., PPC, PFC, 8A). One potential explanation is decreased communication between the areas. Oscillatory coherence is thought to underlie such communication (Fries 2005, Kramer & Eden, 2016; Kass et al., 2014, Singer 1993). Thus, we looked for evidence of altered patterns of LFP coherence during the Unconscious state.

During the Awake state, we found significant stimulus-induced alpha/beta coherence between all pairs of cortical areas for the Cond Tone, the Cond Puff, and the UC Puff (Figure 5B-C, contour lines indicate p < 0.05, Wilcoxon). The UC Tone only induced a small increase in coherence between STG-8A during the Awake state (Figure 5A, contour lines indicate p < 0.05, Wilcoxon). By contrast, during the Unconscious state, there was little or no stimulus-induced coherence for any of the pairs of cortical areas (Figure 5A-C).

**Figure 5:**
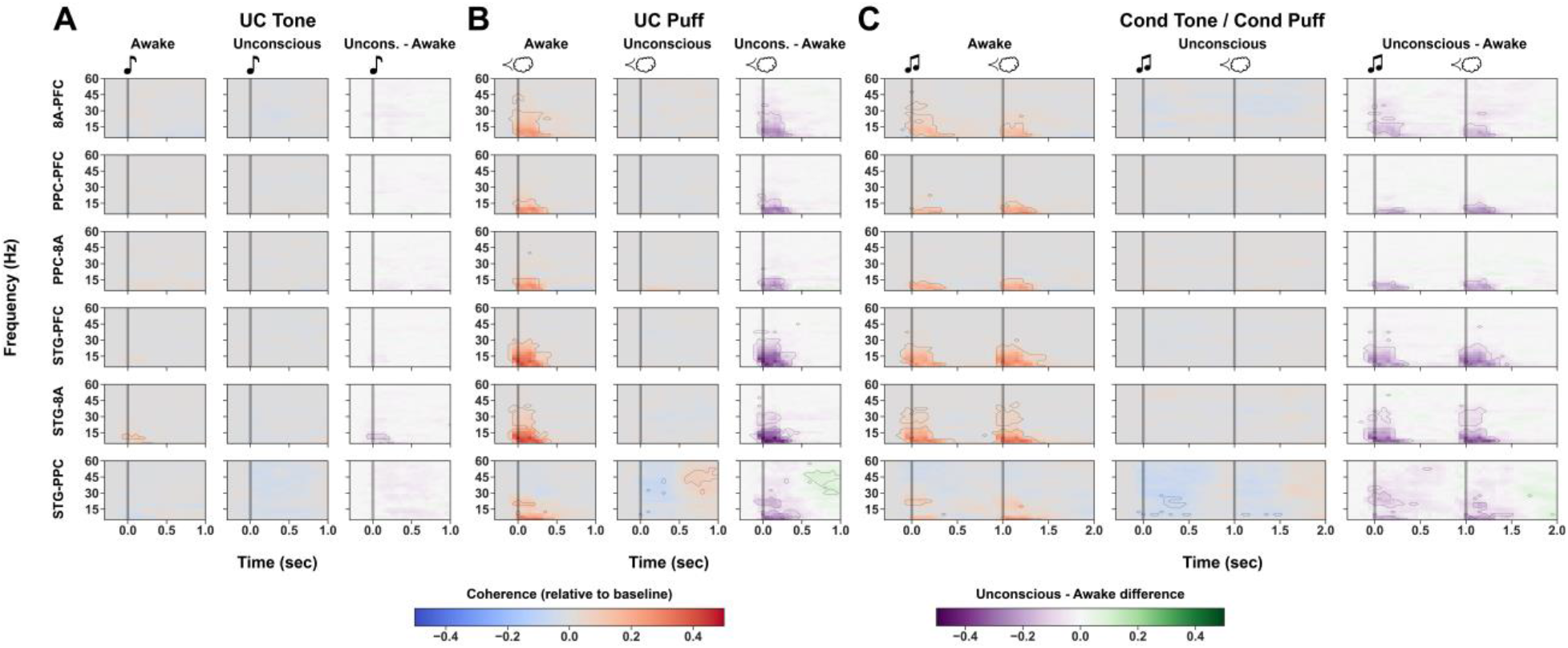
Propofol eliminates stimulus-related coherence between cortical areas. (A-C): Stimulus-evoked change from baseline coherence in Awake (left panel) and Unconscious (middle panel) for indicated stimuli. Right panel shows the difference between Unconscious and Awake. Contour lines show regions of statistical significance (see Methods). Sensory stimuli induced alpha/beta coherence between all cortical areas, which is eliminated by propofol anesthesia.

### Stimulus Responses during Spiking Up states are not Awake-like

Next, we examined the spiking response to sensory stimulation. Previous work has shown propofol anesthesia creates alternating periods of high and low spiking activity referred to, respectively, as Up and Down states (Lewis et al., 2012; Bastos et al., 2021). This raised the possibility that cortical signal transmission might only be impaired during Down states and relatively preserved during Up states. Because standard methods relying on linearity, independence, and identical trials are inadequate in this context, we followed a model-based time-series approach to estimate stimulus responses.

Hidden Markov Models (HMMs) are widely used in statistics and machine learning to model time-series data that alternates between discrete underlying ‘states’ (Rabiner, 1989; Garwood et al., 2021; Chen et al., 2009; Song et al., 2019; Cajigas et al., 2021). We designed and applied an HMM to simultaneously 1) label Up and Down states and 2) estimate population stimulus responses within an area, taking account of the Up and Down states (see Methods for details). While LFP data quality and results were consistent across animals, spike recordings in NHP 2 suffered from complications (see Methods). For this reason, spiking responses for NHP 2 are only reported for PFC.

We found that during the Awake state in NHP 1 there were spiking responses to all stimuli in STG and 8A (Figure 6A-D, shaded regions display 95% CIs). The PPC did not show a spiking response to the UC Tone while the other stimuli elicited an excitatory response. The PFC showed a decrease in spiking in both NHPs to the Cond Tone and the airpuffs, but not the UC Tone. (Figure 6A-D). By contrast, in the Unconscious state responses were drastically reduced. All areas showed minimal spiking to stimuli occurring during Down states. Stimuli that occurred during Up states of STG of NHP 1 evoked responses similar to that observed in the Awake state, but with reduced baseline and maximum firing rates (Figure 6A- D). In PPC and 8A of NHP 1 stimuli occurring in Up states were either absent or dramatically reduced compared to the Awake state (Figure 6A-D). The PFC of both animals did not show spiking responses to stimuli occurring in Up states (Figure 6A-D). Finally, we note that when Up states were aligned across brain areas in NHP 1 no difference was observed in stimulus responses in higher order cortex, but this happened too infrequently to perform a robust analysis (data not shown).

**Figure 6:**
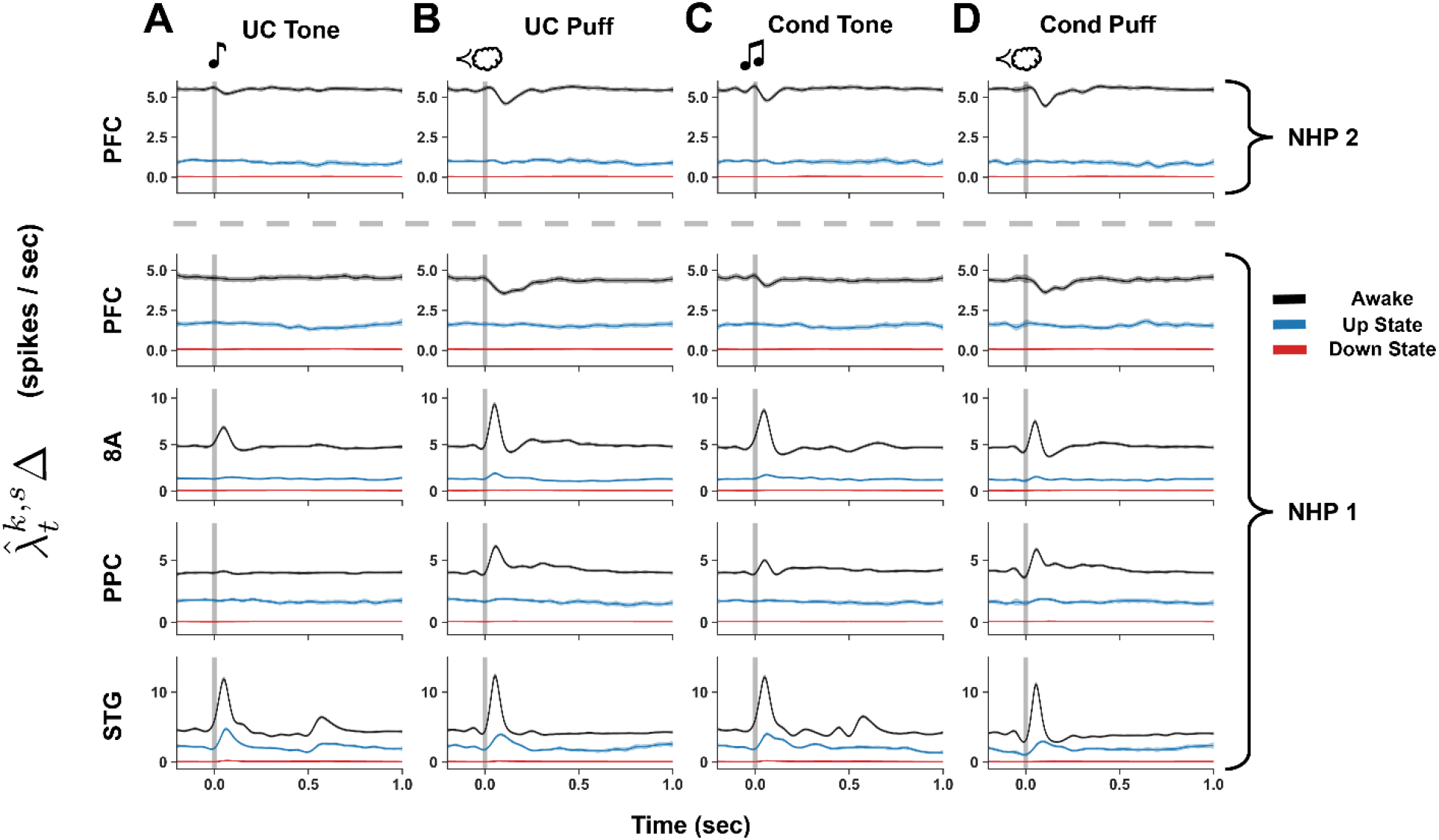
A-D: Estimated population spiking responses to indicated stimulus for Awake (black), Down (red), and Up (blue). For NHP 2, estimated responses are only reported for PFC (see Methods). Shaded regions show 95% confidence intervals.

## Discussion

We found that in the Unconscious state, auditory cortex neural responses to sensory stimuli persisted. However, after auditory cortex, information was progressively lost. In the Awake state, stimulus information was present at all levels of cortex (auditory, parietal, and prefrontal). In the Unconscious state, there was a modest reduction of stimulus information in auditory cortex. In parietal cortex and area 8A (frontal cortex), information was greatly reduced. In prefrontal cortex, stimulus information was reduced to near zero. This could have been due to a breakdown in cortical communication. Stimulus-induced alpha-beta coherence between areas was evident in the Awake state. However, this coherence disappeared in the Unconscious state. Finally, disruption of sensory processing in the Unconscious state consisted of Down states with highly sparse spiking and Up states in which sensory stimulus responses persisted in auditory cortex. Stimulus responses seen in higher-order brain areas in the Awake state were largely lost in Unconscious state.

Many previous studies have examined overall neural state changes due to propofol anesthesia. These changes include an increase in slow-frequency cortical oscillations, a decrease in spiking and high-frequency activity, and a shift of alpha-beta oscillations from posterior to frontal cortex (Purdon et al., 2013; Lewis et al., 2012; Bastos et al., 2021; Redinbaugh et al., 2020; Lewis et al., 2015; Ching et al., 2010; Vijayan et al., 2013). A few studies have examined alterations to sensory stimuli (Krom et al., 2020; Ishizawa et al., 2016; Supp et al., 2011; Liu et al., 2012; Nourski et al., 2017, 2018). They reported results in line with ours, namely a persistence of sensory responses in sensory cortex. Only a few studies have directly compared propofol’s effects on both sensory and higher cortex. A recent study using intracranial electrodes in humans (Krom et al., 2020) found weaker spiking to auditory stimuli in associative cortex but a relative preservation in auditory cortex. A human EEG study (Supp et al., 2011) attributed the loss of sensory responses in higher cortex to increased alpha/beta power. In NHPs, Ishizawa et al. (2016) also found that spiking to sensory inputs persisted in sensorimotor cortex but were reduced in the higher-order premotor area. Our study confirms and expands these observations to additional cortical areas. Under propofol mediated unconsciousness, stimulus-induced inter-areal alpha/beta coherence disappeared. These results suggest propofol may disrupt stimulus-induced coordination of sensory information between cortical areas. The disruption of the alpha/beta bands suggests that top-down feedback may be impaired (Bastos et al., 2015; Van Kerkoerle et al., 2014). Electrical stimulation of the thalamus increases arousal and restored aspects of consciousness (Afrasiabi et al., 2021; Bastos et al., 2021; Redinbaugh et al., 2020). Whether or not it can restore sensory processing will be addressed in future work.

Propofol anesthesia causes an overall shift from higher-frequency to low-frequency activity in cortex. Thus, it was surprising to find that, in auditory cortex, propofol enhanced stimulus-induced broadband power (across all observed frequencies 10-110 Hz). The spectral peak of this effect was in the lower gamma band (30-60 Hz). Given our observation that propofol decreased stimulus-related spiking activity in STG, this increase in broadband power cannot be attributed to an increase in spiking activity (Ray & Maunsell, 2011). A possible explanation is that the increase in stimulus-induced broadband power is due to sub- threshold synaptic effects (Buzsáki et al., 2012; Miller et al., 2014).

Under unconsciousness due to propofol, spiking couples to the phase of the slow wave (0.1 - 4 Hz) (Lewis et al., 2012; Bastos et al., 2021). The results are Down states with little spiking alternating with Up states with increased spiking (Bastos et al., 2021). In theory, propofol-induced disruption of sensory processing could be due only to the Down states. Up states could be small temporal “islands” of normal processing. However, our results did not support that. We found sensory stimuli elicited spiking responses in auditory cortex during Up states that were reduced compared to responses in the Awake state. In higher-order cortical areas, spiking responses observed in the Awake state were dramatically reduced or absent in the Unconscious state. This suggests that Up states do not represent short recapitulations of normal conscious activity but rather are a neurophysiologically distinct state.

Several hypotheses have been proposed to explain disruption of sensory processing during anesthetic induced unconsciousness. The “thalamic gating” hypothesis suggests sensory processing is disrupted at the level of the thalamic relay to primary sensory cortex (Alkire et al., 2000). Our results, along with others (Supp et al., 2011; Boveroux et al., 2010; Raz et al., 2014; Davis et al., 2007; Imas et al., 2005), instead suggest neural responses in sensory cortex, and thus thalamocortical connectivity, are largely preserved. Our results are instead more in line with theories suggesting anesthesia primarily disrupts sensory processing within cortex. One such theory proposes a “multiple hit mechanism”. Many small perturbations add up along sensory processing pathways (Krom et al., 2020). Our results instead suggest a large drop-off between sensory cortex and parietal/frontal cortex. However, we did not record from all of cortex so we may have not observed other perturbations. Our results are also in line with Dehaene and Changeux (2011). They hypothesize conscious awareness relies on broadcasting cortical activity across cortex through long range projections. A disruption of long-range cortical communication would presumably also disrupt the integration of information which the Integrated Information theory suggests is crucial for the conscious state (Tononi et al., 2016).

In sum, we have shown propofol anesthesia eliminates stimulus-related alpha-beta coherence between cortical areas, and progressively disrupts information processing along the cortical hierarchy. These neural signatures of propofol-induced unconsciousness may be fruitful targets for monitoring to avoid intraoperative awareness.

## Data availability

Code used for analysis can be found at github.com/johntauber/propofol_sensory. The data will be made available by reasonable request by contacting the corresponding author.

## Author Contributions

John Tauber: Formal analysis; Methodology; Writing—Original draft; Writing—Review & editing. Scott L. Brincat: Data curation; Investigation; Methodology; Writing—Review & editing. Emily P. Stephen: Methodology; Writing—Review & editing. Jacob A. Donoghue: Investigation. Leo Kozachkov: Software; Writing—Review & editing. Emery N. Brown: Conceptualization; Funding acquisition; Project administration; Supervision; Writing— Review & editing. Earl K. Miller: Conceptualization; Funding acquisition; Project administration; Supervision; Writing—Original draft; Writing—Review & editing.

## Acknowledgements

We thank Andre Bastos, Josefina Correa, Uri Eden, Andrew Eisen and Indie Garwood for helpful comments and feedback.

## Appendix: Additional Technical Details for HMM

To simplify notation moving forward we vectorize the history term component of the CIF as

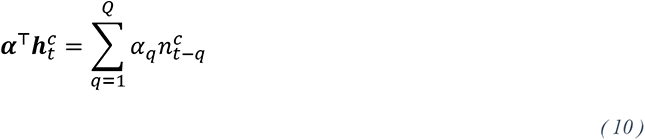

where ***α*** = (*α*_1_, …, *α_Q_*)^⊤^ and 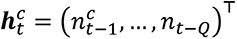. The stimulus terms component is vectorized as

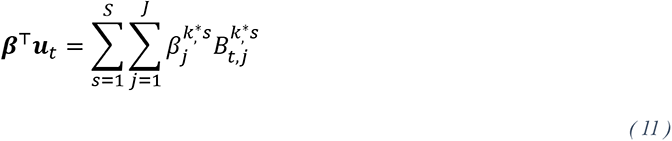

where, using 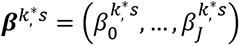 and 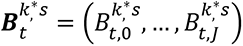 we have concatenated:

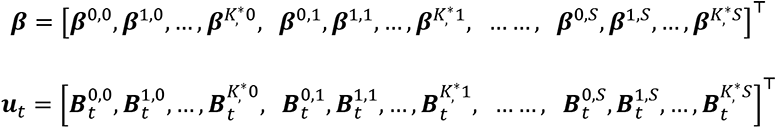

Using this simplified notation, the CIF can now be written:

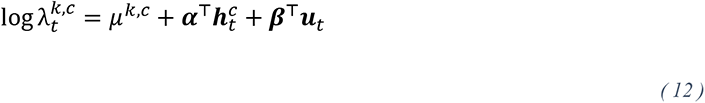

### EM Algorithm

Letting ***n***_***t***_ = (*n_t_*^1^, …, *n_t_*^*C*^)^⊤^, we assume that the neurons are conditionally independent given the hidden state and spiking history and write the joint probability as:

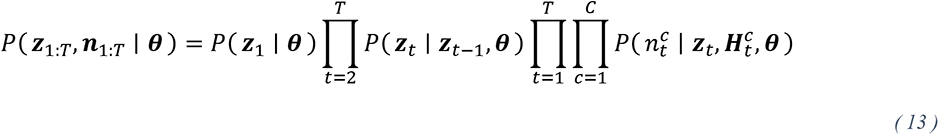

In the EM algorithm, our goal is to maximize the complete-data log likelihood:

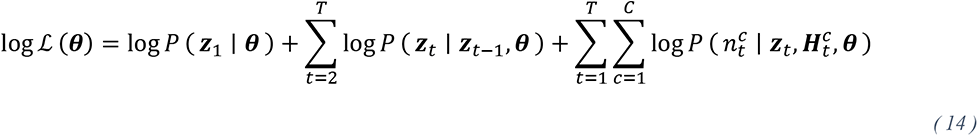

which under our observation model can be written:

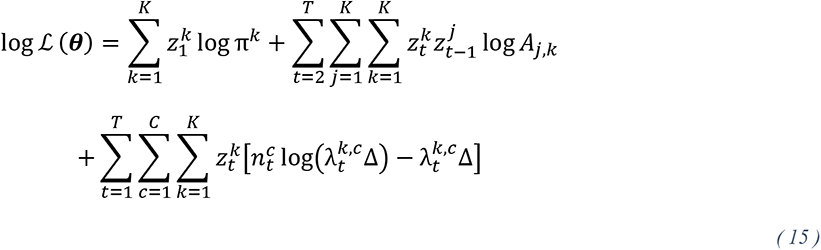

Details on the form of the point process component of (15) can be found in (Smith & Brown, 2003). We define the *Q*-function as the expectation of the complete data log likelihood, on iteration *r* + 1 of the EM algorithm, using the parameters ***θ***^(*r*)^ from the previous iteration:

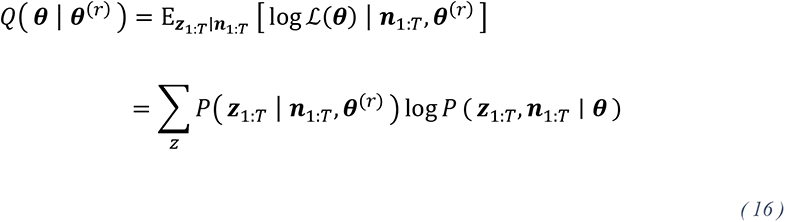

and following (Bishop, 2006) we introduce the following notation:

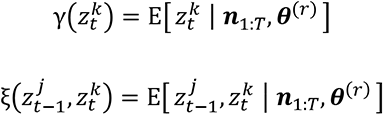

where the expectation is taken over the conditional distribution of the hidden states given the data. We can now write the full *Q*-function:

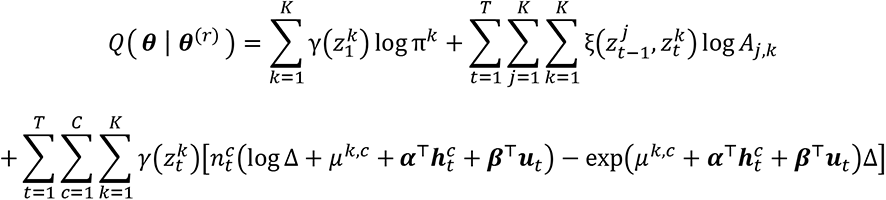

The closed form estimates for ***π*** and **A** are:

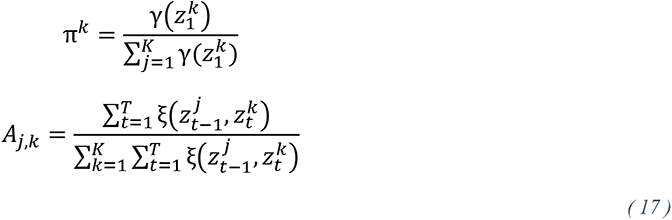

In the BFGS algorithm used to numerically estimate the point-process parameters, we use the following gradients:

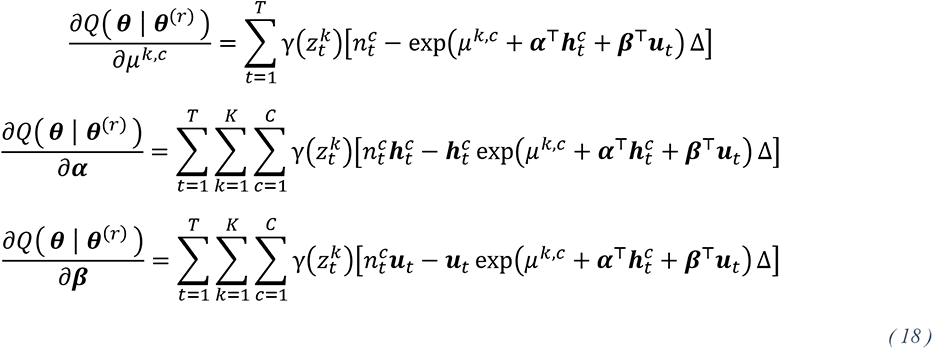

### Louis Method for Confidence Intervals

(Louis, 1982) developed a method which enables calculating the Observed Fisher Information when using the EM algorithm for producing maximum likelihood estimates (Efron & Hinkley, 1978). We use this method for calculating confidence intervals on the estimated spiking response. The method calculates the “observed information” ***I***_***n***_ as:

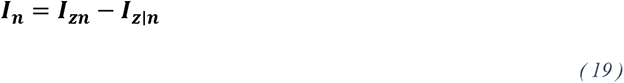

where ***I***_***z***∣***n***_ is the “complete information” calculated from the Hessian of the *Q*-function, and ***I***_***z***∣***n***_ is the “missing information” which provides a correction term to account for the fact that we lose information by *taking the expected value* of the hidden variables.

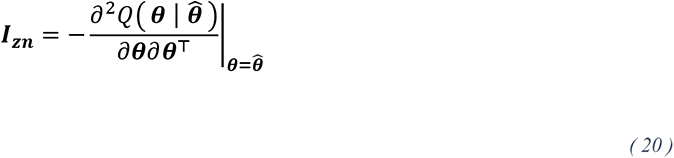

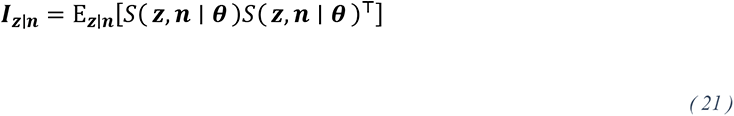

where

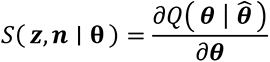

We calculated ***I***_***z***∣***n***_ using a Monte Carlo approximation as in (Turner et al., 1998):

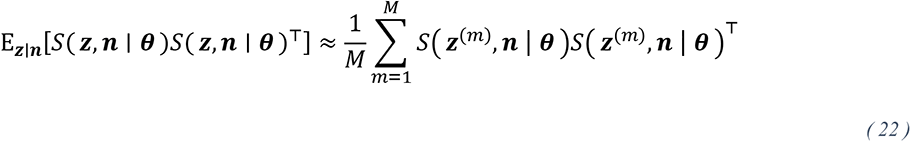

where the ***z***^(*m*)^ are sampled from the posterior distribution of the hidden states given the data. We used *M* = 50 Monte Carlo samples for all sessions. The Hessian of the *Q*-function can be computed using the following:

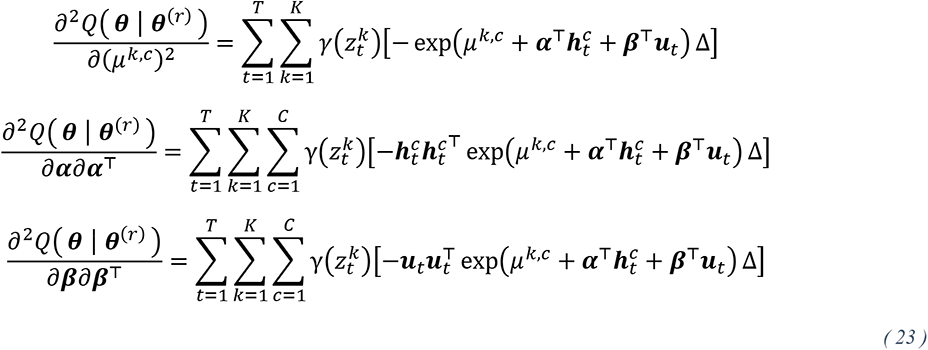

## References

Afrasiabi, M., Redinbaugh, M. J., Phillips, J. M., Kambi, N. A., Mohanta, S., Raz, A., … Saalmann, Y. B. (2021). Consciousness depends on integration between parietal cortex, striatum, and thalamus. Cell Systems, 12 (4), 363–373. doi: 10.1016/j.cels.2021.02.003

Alkire, M. T., Haier, R. J., & Fallon, J. H. (2000). Toward a Unified Theory of Narcosis: Brain Imaging Evidence for a Thalamocortical Switch as the Neurophysiologic Basis of Anesthetic-Induced Unconsciousness. Consciousness and Cognition, 9(3), 370–386. doi: 10.1006/ccog.1999.0423

Babadi, B., & Brown, E. N. (2014). A review of multitaper spectral analysis. IEEE Transactions on Biomedical Engineering, 61(5), 1555–1564. doi: 10.1109/TBME.2014.2311996

Bai, D., Pennefather, P. S., MacDonald, J. F., & Orser, B. A. (1999). The general anesthetic propofol slows deactivation and desensitization of GABA(A) receptors. Journal of Neuroscience, 19(24), 10635–10646. doi: 10.1523/jneurosci.19-24-10635.1999

Bartels, R. H., Beatty, J. C., & Barsky, B. A. (1995). An introduction to splines for use in computer graphics and geometric modeling. Morgan Kaufmann.

Bastos, A. M., Vezoli, J., Bosman, C. A., Schoffelen, J.-M., Oostenveld, R., Dowdall, J. R., . . . Fries, P. (2015). Visual areas exert feedforward and feedback influences through distinct frequency channels. Neuron, 85 (2), 390–401. doi: 10.7554/eLife.60824

Bastos, A. M., Donoghue, J. A., Brincat, S. L., Mahnke, M., Yanar, J., Correa, J., … Miller, E. K. (2021, Apr). Neural effects of propofol-induced unconsciousness and its reversal using thalamic stimulation. eLife, 10, 1–28. doi: 10.7554/eLife.60824

Bishop, C. M. (2006). Pattern recognition and machine learning (Vol. 4) (No. 4). Springer. doi: 10.1117/1.2819119

Boveroux, P., Vanhaudenhuyse, A., Bruno, M. A., Noirhomme, Q., Lauwick, S., Luxen, A., … Boly, M. (2010). Breakdown of within- and between-network resting state functional magnetic resonance imaging connectivity during propofol-induced loss of consciousness. Anesthesiology, 113(5), 1038–1053. doi: 10.1097/ALN.0b013e3181f697f5

Brown, E. N., Lydic, R., & Schiff, N. D. (2010, Dec). General Anesthesia, Sleep, and Coma. New England Journal of Medicine, 363(27), 2638–2650. doi: 10.1056/NEJMra0808281

Brown, E. N., Purdon, P. L., & Van Dort, C. J. (2011, Jul). General Anesthesia and Altered States of Arousal: A Systems Neuroscience Analysis. Annual Review of Neuroscience, 34(1), 601–628. doi: 10.1146/annurev-neuro-060909-153200

Buzsáki, G., Anastassiou, C. A., & Koch, C. (2012). The origin of extracellular fields and currents—EEG, ECoG, LFP and spikes. Nature reviews neuroscience, 13 (6), 407–420. doi: 10.1038/nrn3241

Cajigas, I., Davis, K. C., Meschede-Krasa, B., Prins, N. W., Gallo, S., Naeem, J. A., … others (2021). Implantable brain–computer interface for neuroprosthetic-enabled volitional hand grasp restoration in spinal cord injury. Brain communications, 3(4), doi: 10.1093/braincomms/fcab248

Chen, Z., Vijayan, S., Barbieri, R., Wilson, M. A., & Brown, E. N. (2009). Discrete-and continuous-time probabilistic models and algorithms for inferring neuronal up and down states. Neural computation, 21(7), 1797–1862. doi: 10.1162/neco.2009.06-08-799

Ching, S., Cimenser, A., Purdon, P. L., Brown, E. N., & Kopell, N. J. (2010). Thalamocortical model for a propofol-induced -rhythm associated with loss of consciousness. Proceedings of the National Academy of Sciences, 107(52), 22665–22670. doi: 10.1073/pnas.1017069108

Cramér, H. (1999). Mathematical methods of statistics (Vol. 26). Princeton university press. doi: 10.1515/9781400883868

Crowe, D. A., Goodwin, S. J., Blackman, R. K., Sakellaridi, S., Sponheim, S. R., MacDonald III, A. W., & Chafee, M. V. (2013). Prefrontal neurons transmit signals to parietal neurons that reflect executive control of cognition. Nature Neuroscience, 16 (10), 1484–1491. doi: 10.1038/nn.3509

Daley, D. J., Vere-Jones, D., et al. (2003). An introduction to the theory of point processes: volume i: elementary theory and methods. Springer. doi: 10.1007/b97277

Davis, M. H., Coleman, M. R., Absalom, A. R., Rodd, J. M., Johnsrude, I. S., Matta, B. F., … Menon, D. K. (2007). Dissociating speech perception and comprehension at reduced levels of awareness. Proceedings of the National Academy of Sciences of the United States of America, 104(41), 16032–16037. doi: 10.1073/pnas.0701309104

Dehaene, S., & Changeux, J. P. (2011). Experimental and Theoretical Approaches to Conscious Processing. Neuron, 70(2), 200–227. doi: 10.1016/j.neuron.2011.03.018

Dempster, A. P., Laird, N. M., & Rubin, D. B. (1977). Maximum likelihood from incomplete data via the EM algorithm. Journal of the Royal Statistical Society. Series B (Methodological), 39(1), 1–38. doi: 10.1111/j.2517-6161.1977.tb01600.x

Efron, B., & Hinkley, D. V. (1978, Dec). Assessing the accuracy of the maximum likelihood estimator: Observed versus expected Fisher information. Biometrika, 65(3), 457–483. doi: 10.1093/biomet/65.3.457

Fletcher, R. (2013). Practical methods of optimization. John Wiley & Sons. doi: 10.1002/9781118723203

Fries, P. (2005). A mechanism for cognitive dynamics: neuronal communication through neuronal coherence. Trends in cognitive sciences, 9(10), 474–480. doi: 10.1016/j.tics.2005.08.011

Fries, P. (2015, Oct). Rhythms for Cognition: Communication through Coherence. Neuron, 88(1), 220–235. doi: 10.1016/j.neuron.2015.09.034

Garwood, I. C., Chakravarty, S., Donoghue, J., Mahnke, M., Kahali, P., Chamadia, S., … Brown, E. N. (2021). A hidden markov model reliably characterizes ketamine-induced spectral dynamics in macaque local field potentials and human electroencephalograms. PLoS Computational Biology, 17(8), e1009280. doi: 10.1371/journal.pcbi.1009280

Ghoneim, M. M. (2000). Awareness during Anesthesia. Anesthesiology, 92(2), 597–602. doi: 10.1213/ane.0b013e318193c634

Hamilton, J. D. (1990). Analysis of time series subject to changes in regime. Journal of Econometrics, 45(1- 2), 39–70. doi: 10.1016/0304-4076(90)90093-9

Hastie, T., Tibshirani, R., Friedman, J. H., & Friedman, J. H. (2009). The elements of statistical learning: data mining, inference, and prediction (Vol. 2). Springer. doi: 10.1007/978-0-387-84858-7

Hemmings, H. C., Akabas, M. H., Goldstein, P. A., Trudell, J. R., Orser, B. A., & Harrison, N. L. (2005). Emerging molecular mechanisms of general anesthetic action. Trends in Pharmacological Sciences, 26(10), 503–510. doi: 10.1016/j.tips.2005.08.006

Hemmings, H. C., Riegelhaupt, P. M., Kelz, M. B., Solt, K., Eckenhoff, R. G., Orser, B. A., & Goldstein, P. A. (2019). Towards a Comprehensive Understanding of Anesthetic Mechanisms of Action: A Decade of Discovery. Trends in Pharmacological Sciences, 40(7), 464–481. doi: 10.1016/j.tips.2019.05.001

Imas, O. A., Ropella, K. M., Ward, B. D., Wood, J. D., & Hudetz, A. G. (2005). Volatile anesthetics disrupt frontal-posterior recurrent information transfer at gamma frequencies in rat. Neuroscience Letters, 387(3), 145–150. doi: 10.1016/j.neulet.2005.06.018

Ishizawa, Y., Ahmed, O. J., Patel, S. R., Gale, J. T., Sierra-Mercado, D., Brown, E. N., & Eskandar, E. N. (2016). Dynamics of propofol-induced loss of consciousness across primate neocortex. Journal of Neuroscience, 36(29), 7718–7726. doi: 10.1523/JNEUROSCI.4577-15.2016

Jarvis, M., & Mitra, P. P. (2001). Sampling properties of the spectrum and coherency of sequences of action potentials. Neural Computation, 13 (4), 717–749. doi: 10.1162/089976601300014312

Kass, R. E., Eden, U. T., & Brown, E. N. (2014). Analysis of neural data (Vol. 491). Springer. doi: 10.1007/978-1-4614-9602-1

Kotsovolis, G., & Komninos, G. (2009). Awareness during anesthesia: How sure can we be that the patient is sleeping indeed? Hippokratia, 13(2), 83–89.

Kramer, M. A., & Eden, U. T. (2016). Case studies in neural data analysis: a guide for the practicing neuroscientist. MIT Press.

Krom, A. J., Marmelshtein, A., Gelbard-Sagiv, H., Tankus, A., Hayat, H., Hayat, D., … Nir, Y. (2020). Anesthesia-induced loss of consciousness disrupts auditory responses beyond primary cortex. Proceedings of the National Academy of Sciences, 201917251. doi: 10.1073/pnas.1917251117

Lewis, L. D., Ching, S., Weiner, V. S., Peterfreund, R. A., Eskandar, E. N., Cash, S. S., … Purdon, P. (2013). Local cortical dynamics of burst suppression in the anaesthetized brain. Brain, 136, 2727–2737. doi: 10.1093/brain/awt174

Lewis, L. D., Voigts, J., Flores, F. J., Ian Schmitt, L., Wilson, M. A., Halassa, M. M., & Brown, E. N. (2015). Thalamic reticular nucleus induces fast and local modulation of arousal state. eLife, 4(OCTOBER2015), 1–23. doi: 10.7554/eLife.08760

Lewis, L. D., Weiner, V. S., Mukamel, E. A., Donoghue, J. A., Eskandar, E. N., Madsen, J. R., … Purdon, P. L. (2012, Dec). Rapid fragmentation of neuronal networks at the onset of propofol-induced unconsciousness. Proceedings of the National Academy of Sciences, 109(49), E3377–E3386. doi: 10.1073/pnas.1210907109

Linderman, S., Antin, B., Zoltowski, D., & Glaser, J. (2020). SSM: Bayesian Learning and Inference for State Space Models. Retrieved from https://github.com/lindermanlab/ssm

Liu, X., Lauer, K. K., Ward, B. D., Rao, S. M., Li, S. J., & Hudetz, A. G. (2012). Propofol disrupts functional interactions between sensory and high-order processing of auditory verbal memory. Human Brain Mapping, 33(10), 2487–2498. doi: 10.1002/hbm.21385

Louis, T. A. (1982). Finding the observed information matrix when using the em algorithm. Journal of the Royal Statistical Society. Series B (Methodological), 44(2), 226–233. doi: 10.1111/j.2517-6161.1982.tb01203.x

Miller, K. J., Honey, C. J., Hermes, D., Rao, R. P., Ojemann, J. G., et al. (2014). Broadband changes in the cortical surface potential track activation of functionally diverse neuronal populations. Neuroimage, 85, 711–720.

Murphy, K. P. (2023). Probabilistic machine learning: Advanced topics. MIT Press. doi: 10.1016/j.neuroimage.2013.08.070

Nourski, K. V., Banks, M. I., Steinschneider, M., Rhone, A. E., Kawasaki, H., Mueller, R. N., … Howard, M. A. (2017). Electrocorticographic delineation of human auditory cortical fields based on effects of propofol anesthesia. NeuroImage, 152(February), 78–93. doi: 10.1016/j.neuroimage.2017.02.061

Nourski, K. V., Steinschneider, M., Rhone, A. E., Kawasaki, H., Howard, M. A., & Banks, M. I. (2018). Auditory predictive coding across awareness states under anesthesia: An intracranial electrophysiology study. Journal of Neuroscience, 38(39), 8441–8452. doi: 10.1523/JNEUROSCI.0967-18.2018

Pawitan, Y. (2001). In all likelihood: statistical modelling and inference using likelihood. Oxford University Press.

Pedregosa, F., Varoquaux, G., Gramfort, A., Michel, V., Thirion, B., Grisel, O., … others (2011). Scikit-learn: Machine learning in python. Journal of machine learning research, 12(Oct), 2825–2830. doi: 10.5555/1953048.2078195

Pesaran, B., Vinck, M., Einevoll, G. T., Sirota, A., Fries, P., Siegel, M., … Srinivasan, R. (2018, Jul). Investigating large-scale brain dynamics using field potential recordings: analysis and interpretation. Nature Neuroscience, 21(7), 903–919. doi: 10.1038/s41593-018-0171-8

Purdon, P. L., Pierce, E. T., Mukamel, E.A., Prerau, M. J., Walsh, J. L., Wong, K. F. K., … Brown, E. N. (2013, Mar). Electroencephalogram signatures of loss and recovery of consciousness from propofol. Proceedings of the National Academy of Sciences, 110(12), E1142–E1151. doi: 10.1073/pnas.1221180110

Purdon, P. L., Sampson, A., Pavone, K. J., & Brown, E. N. (2015, Oct). Clinical Electroencephalography for Anesthesiologists. Anesthesiology, 123(4), 937–960. doi: 10.1097/ALN.0000000000000841

Rabiner, L. R. (1989). A tutorial on hidden markov models and selected applications in speech recognition. Proc. IEEE, 77, 257–286. doi: 10.1109/5.18626

Ray, S., & Maunsell, J. H. (2011). Different origins of gamma rhythm and high-gamma activity in macaque visual cortex. PLoS Biology, 9(4). doi: 10.1371/journal.pbio.1000610

Raz, A., Grady, S. M., Krause, B. M., Uhlrich, D. J., Manning, K.A., & Banks, M. I. (2014). Preferential effect of isoflurane on top-down vs. bottom-up pathways in sensory cortex. Frontiers in Systems Neuroscience, 8(OCT), 1–22. doi: 10.3389/fnsys.2014.00191

Redinbaugh, M. J., Phillips, J. M., Kambi, N. A., Mohanta, S., Andryk, S., Dooley, G. L., … Saalmann, Y. B. (2020). Thalamus Modulates Consciousness via Layer-Specific Control of Cortex. Neuron, 106(1), 66–75.e12. doi: 10.1016/j.neuron.2020.01.005

Sahinovic, M. M., Struys, M. M., & Absalom, A. R. (2018). Clinical Pharmacokinetics and Pharmacodynamics of Propofol. Clinical Pharmacokinetics, 57(12), 1539–1558. doi: 10.1007/s40262-018-0672-3

Seabold, S., & Perktold, J. (2010). Statsmodels: Econometric and statistical modeling with python. In 9th Python in Science Conference. doi: 10.25080/Majora-92bf1922-011

Sebel, P. S., Bowdle, T. A., Ghoneim, M. M., Rampil, I. J., Padilla, R. E., Gan, T. J., & Domino, K. B. (2004). The incidence of awareness during anesthesia: A multicenter United States study. Anesthesia and Analgesia, 99(3), 833–839. doi: 10.1213/01.ANE.0000130261.90896.6C

Singer, W. (1993). Synchronization of cortical activity and its putative role in information processing and learning. Annual review of physiology, 55(1), 349–374. doi: 10.1146/annurev.ph.55.030193.002025

Smith, A. C., & Brown, E. N. (2003, May). Estimating a State-Space Model from Point Process Observations. Neural Computation, 15(5), 965–991. doi: 10.1162/089976603765202622

Song, A. H., Chlon, L., Soulat, H., Tauber, J., Subramanian, S., Ba, D., & Prerau, M. J. (2019). Multitaper infinite hidden markov model for EEG. In 2019 41st Annual International Conference of the IEEE Engineering in Medicine and Biology Society (EMBC) (pp. 5803–5807). doi: 10.1109/EMBC.2019.8856817

Supp, G. G., Siegel, M., Hipp, J. F., & Engel, A. K. (2011). Cortical Hypersynchrony Predicts Breakdown of Sensory Processing during Loss of Consciousness. Current Biology, 21, 1988–1993. doi: 10.1016/j.cub.2011.10.017

Tononi, G., Boly, M., Massimini, M., & Koch, C. (2016). Integrated information theory: from consciousness to its physical substrate. Nature Reviews Neuroscience, 17(7), 450–461. doi: 10.1038/nrn.2016.44

Turner, T. R., Cameron, M. A., & Thomson, P. J. (1998). Hidden markov chains in generalized linear models. The Canadian Journal of Statistics / La Revue Canadienne de Statistique, 26(1), 107–125. doi: 10.2307/3315677

Van Kerkoerle, T., Self, M. W., Dagnino, B., Gariel-Mathis, M.-A., Poort, J., Van Der Togt, C., & Roelfsema, P. R. (2014). Alpha and gamma oscillations characterize feedback and feedforward processing in monkey visual cortex. Proceedings of the National Academy of Sciences, 111 (40), 14332–14341. doi: 10.1073/pnas.1402773111

Vijayan, S., Ching, S. N., Purdon, P. L., Brown, E. N., & Kopell, N. J. (2013). Thalamocortical mechanisms for the anteriorization of alpha rhythms during propofol-induced unconsciousness. Journal of Neuroscience, 33(27), 11070–11075. doi: 10.1523/JNEUROSCI.5670-12.2013

Virtanen, P., Gommers, R., Oliphant, T. E., Haberland, M., Reddy, T., Cournapeau, D., … others (2020). Scipy 1.0: fundamental algorithms for scientific computing in python. Nature methods, 17(3), 261–272. doi: 10.1038/s41592-019-0686-2

